# Low copy CRISPR-Cas13d mitigates collateral RNA cleavage

**DOI:** 10.1101/2024.05.13.594039

**Authors:** Sydney K. Hart, Hans-Hermann Wessels, Alejandro Méndez-Mancilla, Simon Müller, Gediminas Drabavicius, Olivia Choi, Neville E. Sanjana

## Abstract

While CRISPR-Cas13 systems excel in accurately targeting RNA, the potential for collateral RNA degradation poses a concern for therapeutic applications and limits broader adoption for transcriptome perturbations. We evaluate the extent to which collateral RNA cleavage occurs when *Rfx*Cas13d is delivered via plasmid transfection or lentiviral transduction and find that collateral activity only occurs with high levels of *Rfx*Cas13d expression. Using transcriptome-scale and combinatorial CRISPR pooled screens in cell lines with low-copy *Rfx*Cas13d, we find high on-target knockdown, without extensive collateral activity regardless of the expression level of the target gene. In contrast, transfection of *Rfx*Cas13d, which yields higher nuclease expression, results in collateral RNA degradation. Further, our analysis of a high-fidelity Cas13 variant uncovers a marked decrease in on-target efficiency, suggesting that its reduced collateral activity may be due to an overall diminished nuclease capability.

## Main

Type VI CRISPR-Cas nucleases, and specifically *Rfx*Cas13d, have been harnessed for a variety of RNA-targeting applications due to their precise and efficient knockdown of RNA transcripts^1–3^. RNA-targeting CRISPR systems have demonstrated therapeutic potential for temporary or non-heritable edits^2,3^; however, collateral RNA cleavage in Cas13-based RNA editing systems may limit their use for transcriptome editing *in vivo*. Although initial studies using *Rfx*Cas13d *in vitro* and *in vivo* did not detect collateral RNA degradation^1,4,5^, several recent studies have demonstrated collateral activity when highly-expressed genes are targeted.^6–9^ Resolving these discrepancies requires a systematic characterization of the extent and context of collateral RNA targeting. A thorough understanding of this phenomenon and methods to mitigate it would enhance the utility of Cas13 for RNA-targeting assays, transcriptome-scale screens of coding and noncoding RNAs, and the safety of future Cas13-based therapies.

One initial observation we made was that studies employing transfection methods such as piggyBac transposons^8,10^ or liposome-mediated transfection^6–8^ of *Rfx*Cas13d reported higher frequencies of collateral activity than studies using viral delivery of similar constructs^1,3–5,11–13^. Consistent with previous work suggesting that regulating *Rfx*Cas13d expression reduces collateral activity^6^, we hypothesized that the disparity in reported collateral activity between studies using viral or transfection-based delivery of *Rfx*Cas13d might be due to differences in *Rfx*Cas13d expression. We found that lentiviral transduction results in lower levels of transgene expression than plasmid transfection, either when directly quantifying *Rfx*Cas13d expression (**Fig. 1a, Supplementary Fig. 1a,b**) or when using a reporter gene on the same vector (**Supplementary Fig. 1c-e**).

**Figure 1.**
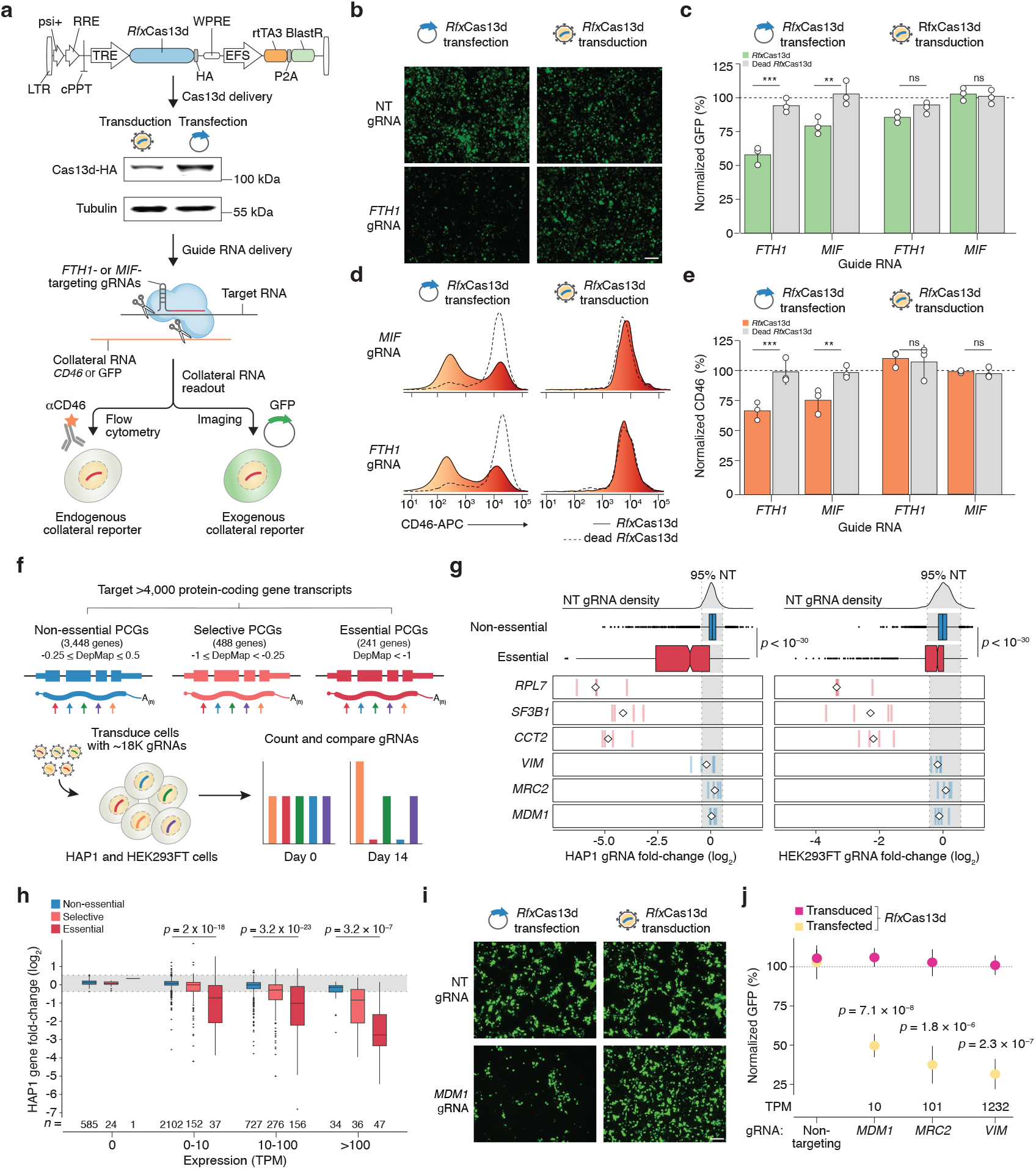
Collateral RNA cleavage occurs with transfection but not transduction of *Rfx*Cas13d. **a**, Collateral RNA cleavage assay to compare transfection- and transduction-based delivery of *Rfx*Cas13d using a GFP reporter or endogenous CD46 expression. A doxycycline-inducible Cas13d vector was either transfected or transduced into HEK293FT cells. In both cases, cells were transfected with Cas13 guide RNAs (gRNAs) targeting *FTH1* or *MIF*. Western blot depicts HA-tagged *Rfx*Cas13d protein expression. **b**, Representative images of human HEK293FT cells co-transfected with GFP and *FTH1*-targeting or non-targeting (NT) gRNAs. Cells were either transduced or transfected with Cas13d. Scale bar: 100 μm. **c**, GFP expression after co-transfection of a GFP reporter and gRNAs targeting *FTH1* or *MIF* in HEK293FT cells. Prior to this co-transfection, cells were either transduced or transfected with active or catalytically dead Cas13d. GFP expression was normalized to non-targeting control gRNAs and significance was determined using a two-sided *t*-test (*n* = 3 biological replicates): ** *p* < 10^−2^, *** *p* < 10^−3^, ns, not significant. **d**, Representative cell surface expression of CD46 in HEK293FT cells transduced or transfected with Cas13d (either active or a catalytically inactive variant) after transfection of gRNAs targeting *FTH1* or *MIF* in HEK293FT cells. **e**, CD46 expression in HEK293FT cells transfected with gRNAs targeting *FTH1* or *MIF*. Prior to this co-transfection, cells were either transduced or transfected with active or catalytically dead Cas13d. CD46 expression was normalized to non-targeting control gRNAs and significance was determined using a two-sided *t*-test (*n* = 3 biological replicates): ** *p* < 10^−2^, *** *p* < 10^−3^, ns, not significant. **f**, A Cas13d pooled screen with 17,708 gRNAs targeting essential and nonessential protein-coding genes (and non-targeting controls) with different transcript expression levels (*n* = 4,177 genes covering 20% of the human protein-coding transcriptome) in HEK293FT and HAP1 cells (*n* = 2 biological replicate screens per cell line). After 14 days of culture, we identified whether gRNAs targeting essential, selective, and nonessential genes were depleted by comparing abundance between day 14 and day 0. **g**, Normalized depletion of individual gRNAs targeting essential (*red*) and nonessential (*blue*) genes (*n* = 4 gRNAs). The median of the gRNAs is indicated by the diamond. The distribution of the middle 95% of non-targeting (NT) gRNAs is shown in *gray*. Boxplots indicate all gRNAs targeting essential (mean DepMap Chronos score < -1, *n* = 1,095 cell lines) (*red*) and nonessential (Chronos > -0.25) (*blue*) genes in HEK293FT and HAP1 cells and significance was determined using a two-sided Mann-Whitney *U* test. **h**, Comparison of fold-change and expression level of essential, selective, and nonessential genes in HAP1 cells. Dashed lines indicate the distribution of the middle 95% of NT gRNAs. Statistical significance between nonessential and essential gene categories was determined by a two-sided Mann-Whitney *U* test. **i**, Representative images of human HEK293FT cells co-transfected with GFP and *MDM1*-targeting or non-targeting (NT) gRNAs. *MDM1* is a nonessential gene, as indicated in panel *g*. Cells were either transduced or transfected with Cas13d. Scale bar: 100 μm. **j**, GFP expression after co-transfection of a GFP reporter vector and gRNAs targeting nonessential genes *MDM1, MRC2*, and *VIM* in HEK293FT cells. Prior to this co-transfection, cells were either transduced or transfected with active or catalytically dead Cas13d. GFP expression was normalized to cells with a catalytically dead Cas13d (transduced or transfected, as indicated) and significance was determined using two-sided *t*-test (mean of 2 gRNAs, *n* = 3 biological replicates).

To determine if these differences in *Rfx*Cas13d expression led to collateral activity, we compared the levels of collateral RNA degradation using either plasmid transfection or lentiviral transduction of *Rfx*Cas13d while targeting the highly-expressed genes *FTH1* and *MIF* (**Fig. 1a**); these genes were previously reported by others to lead to collateral activity using a fluorescent reporter^7^, although DepMap^14^ indicates that *FTH1* may be selectively essential (**Supplementary Fig. 1f**). Using a similar readout for collateral activity, we found that transfection of *Rfx*Cas13d leads to a reduction in GFP protein via quantitative microscopy (**Fig. 1b, c**) and that this collateral cleavage is dose-dependent (**Supplementary Fig. 1g, h**). In contrast, lentiviral transduction at a low multiplicity-of-infection (< 0.3) produced no difference in GFP expression compared to non-targeting controls (**Fig. 1b, c**).

We next sought to understand whether similar collateral activity occurs with endogenous transcripts. Using flow cytometry, we quantified expression of CD46, a ubiquitously-expressed cell surface protein, when targeting *FTH1* and *MIF*. Consistent with our GFP readout, we found that lentiviral transduction of *Rfx*Cas13d did not result in the collateral RNA degradation of *CD46* but transfection did (**Fig. 1d, e**) — even though on-target knockdown was similar in both settings (**Supplementary Fig. 1i-k**).

Given the absence of collateral activity on exogenous and native transcripts when targeting these two genes (*FTH1* and *MIF*) via viral transduction, we sought to assess this more comprehensively across the transcriptome: We performed a pooled Cas13 screen to target ∼20% of all protein-coding genes covering a diverse range of gene expression levels in two human cell lines (HAP1 and HEK293FT) (**Fig. 1f, Supplementary Fig. 2a, Supplementary Table 1**). In total, we targeted 3,448 nonessential genes (as classified by DepMap^14^) with four gRNAs each. As positive controls for depletion, we also included 241 essential genes and 488 selective (essential but not strongly so) genes. Previous studies of Cas13 collateral activity have shown a substantial reduction in cell viability, which has been attributed to RNase-like chromatin collapse^7^ and cleavage of 28S ribosomal RNAs^7,9^. Thus, we reasoned that, if collateral activity occurs, we would observe a depletion of nonessential but highly-expressed genes.

**Figure 2.**
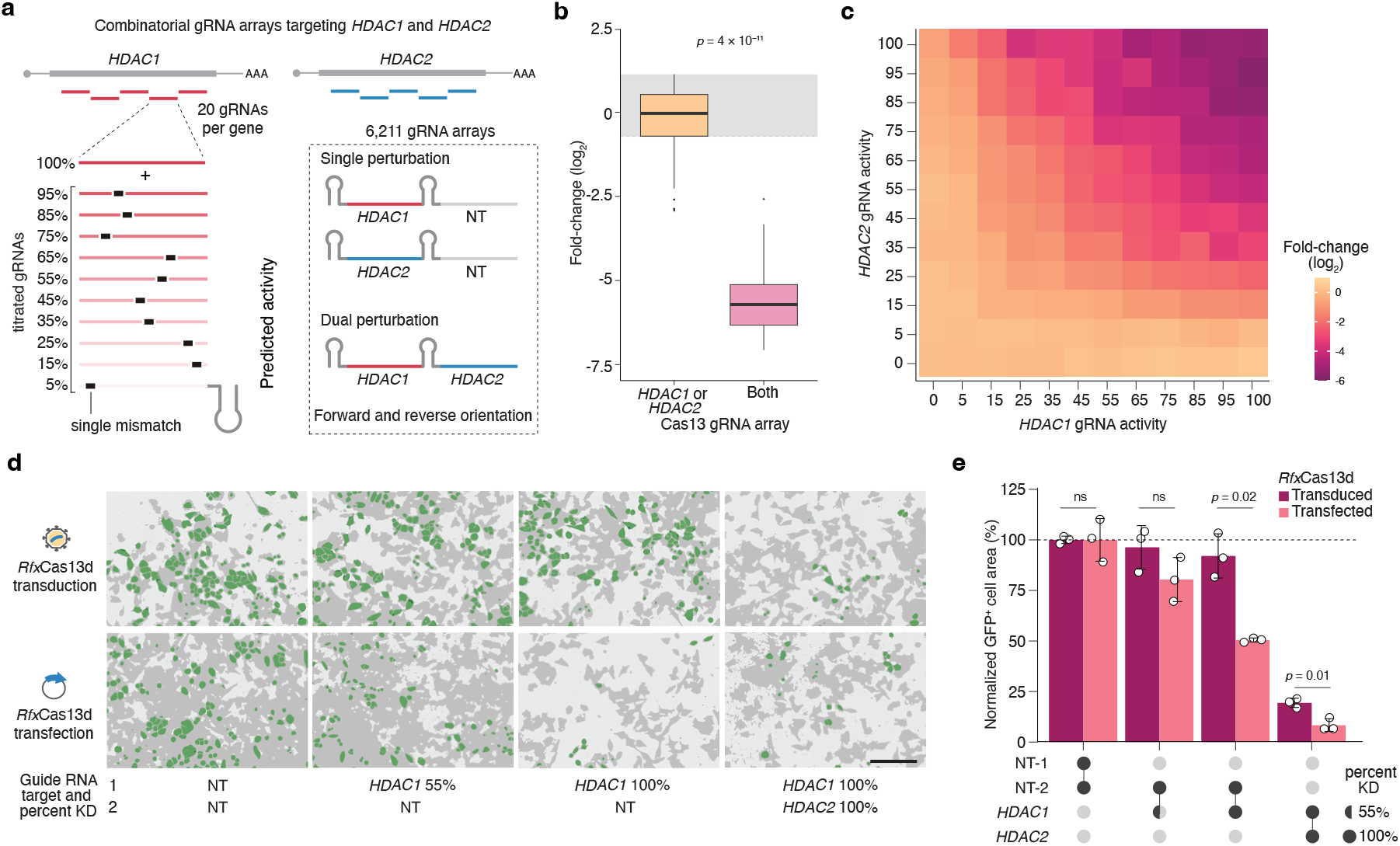
Combinatorial knockdown of synthetic lethal pair *HDAC1* - *HDAC2* confirms lack of collateral RNA cleavage after low copy-number transduction of *Rfx*Cas13d. **a**, A combinatorial titration pooled transcriptomic screen targeting a synthetic lethal gene pair (*HDAC1* and *HDAC2*). Designed position-specific mismatches in the gRNA arrays titrate knockdown activity of *Rfx*Cas13d. After gRNA library transduction into human A375 cells, depletion of single or dual gene perturbations is quantified by comparing gRNA counts between day 14 and day 0. **b**, Fold-change of gRNAs targeting *HDAC1* or *HDAC2* individually or together at 100% predicted gRNA activity 14 days after *Rfx*Cas13d induction (*n* = 120 gRNA arrays with 40 each for *HDAC1* only, *HDAC2* only and both *HDAC1* and *HDAC2* targeting). The distribution of the central 95% of non-targeting gRNAs is shown in *gray* (*n* = 492 non-targeting perturbations). Statistical significance determined by two-sided Mann-Whitney *U* test. **c**, Median fold-change of dual gRNA perturbations (targeting *HDAC1* and *HDAC2*), binned based on predicted gRNA activity (*n* = 40 gRNA arrays per gene-targeting heatmap tile, *n* = 492 gRNA arrays for non-targeting perturbations [0 *x* 0 heatmap tile]). **d**, Representative images of a competition assay using A375 cells transduced with dual gRNA arrays targeting *HDAC1* and *HDAC2* at 55% or 100% predicted gRNA activity or non-targeting gRNAs and imaged 5 days later. The dual gRNA array constructs also constitutively express GFP; these cells were mixed 1:1 with cells without a dual gRNA array (non-fluorescent cells) at the time of plating. Cells with and without GFP/array were either transduced or transfected with *Rfx*Cas13d as indicated. Scale bar: 200 μm. **e**, Percent of GFP-positive cells for each gRNA array normalized to non-targeting gRNA array control for both transduced and transfected *Rfx*Cas13d cells (*n* = 27 images from 3 independent transductions/transfections). Statistical significance determined with a two-sided *t*-test. ns, not significant.

In line with our prior experiments targeting *FTH1* and *MIF*, we found that nonessential genes do not deplete in the pooled screen regardless of their expression levels. In contrast, essential genes readily deplete from the screen, demonstrating that on-target knockdown is highly efficient (**Fig. 1g, h, Supplementary Fig. 2b, c, Supplementary Table 2**). As expected, selective genes from DepMap also deplete but not to the same degree as those classified as essential. Subsequently, we conducted additional validation of three nonessential genes from this screen with low (*MDM1*, 10 transcripts per million [TPM]), medium (*MRC2*, 101 TPM), and high (*VIM*, 1232 TPM) expression, respectively, and did so in HEK293FT cells that were either transduced with *Rfx*Cas13d lentivirus or transfected with the same vector. Again, using co-transfected GFP as a readout for collateral activity, we found that only transfection of *Rfx*Cas13d results in collateral GFP degradation and that GFP depletion positively correlates with the targeted gene’s expression (**Fig. 1i, j, Supplementary Fig. 2d**).

Next, we investigated whether modifying the guide RNA sequence could diminish the recognition of the target RNA transcript and hinder the conformational changes necessary for *cis* and *trans* RNA cleavage. We performed a combinatorial *Rfx*Cas13d screen in A375 cells targeting *HDAC1* and/or *HDAC2* using ∼6,000 pairs of gRNAs. These two genes form a synthetic lethal gene pair, where targeting both genes (but not either singly) results in lethality and drop-out, as shown in a recent study using DNA-targeting perturbations^15^. We took advantage of our TIGER (Targeted Inhibition of Gene Expression via gRNA design) model^13^ to design titrated gRNAs with single mismatches in the parent sequence that alter the expected transcript knockdown activity over a range of partial knockdowns (**Fig. 2a, Supplementary Fig. 3a, Supplementary Table 3**). Targeting a single gene from the pair should decrease cell fitness due to the high expression levels of these genes —147 TPM for *HDAC1* and 130 TPM for *HDAC2* — if unintended collateral activity occurs.

**Figure 3.**
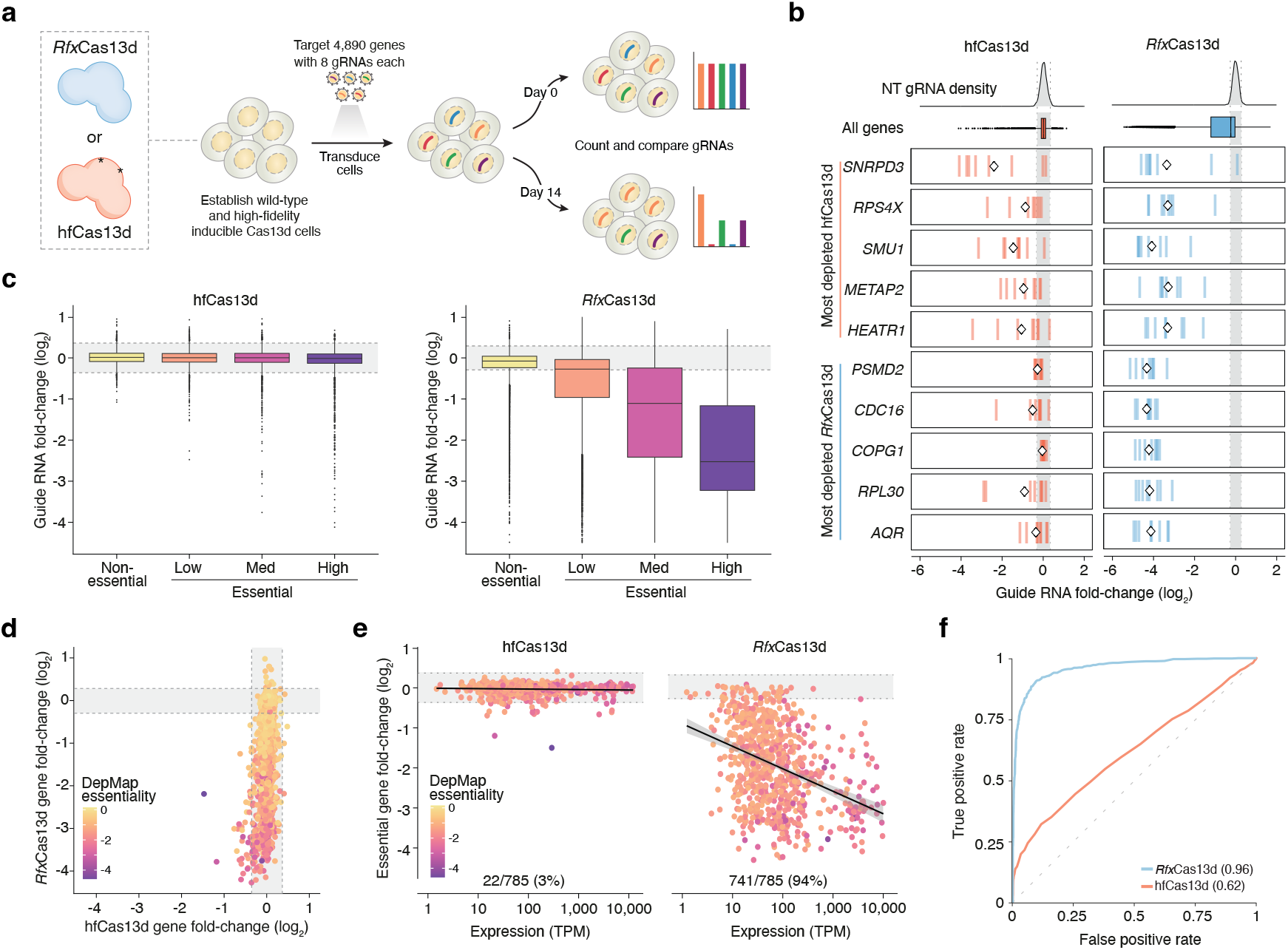
A high-fidelity Cas13d variant (hfCas13d) that minimizes collateral RNA cleavage has substantially less on-target knockdown. **a**, The *Rfx*Cas13d and hfCas13d screens compare nuclease-specific differences in gene depletion for low copy-number transduction. After library transduction into HAP1 cells, gRNA depletion is quantified by comparing gRNA counts between day 14 and day 0. **b**, Depletion of gRNAs targeting the top five most depleted genes from the hfCas13d (*orange*) and *Rfx*Cas13d (*blue*) screens (*n* = 8 gRNAs per gene). The mean of the eight gRNAs is indicated by the diamond. The densities of non-targeting (NT) gRNAs and all gene-targeting gRNAs are also shown. **c**, For the indicated Cas13 nuclease, fold-change of gRNAs, categorized by DepMap gene essentiality (DepMap release 05-2023)^14^: Nonessential (mean DepMap Chronos score ≥ -0.1), low (−0.5 < Chronos < - 0.1), medium (−1 ≤ Chronos ≤ -0.5), and high (Chronos < -1) essentiality (*n* = 1,095 cell lines). **d**, Fold change of all genes in the *Rfx*Cas13d and hfCas13d screens (*n* = 4,890 genes). Color denotes gene essentiality (mean DepMap Chronos score). **e**, Depletion of highly-essential genes (mean DepMap Chronos score < -1) as a function of target gene expression genes (*n* = 785 highly-essential genes). Each gene is colored by its mean Chronos score. The percent of genes depleted is quantified based on whether a gene is more depleted than the 99th percentile of the non-targeting gRNA distribution. **f**, Receiver-operator characteristic (ROC) curve for DepMap highly-essential genes compared to nonessential genes (*n* = 785 highly-essential genes, *n* = 1,985 nonessential genes). Numbers in parenthesis next to each Cas13 protein indicate area under the ROC (AUROC) curve.

Consistent with our previous data, we found that targeting only one gene, *HDAC1* or *HDAC2*, from this synthetic lethal gene pair does not cause significant gRNA depletion in our lentiviral, low-copy *Rfx*Cas13d cell lines (**Fig. 2b, Supplementary Fig. 3b, Supplementary Fig. 4a, b**). As expected, targeting both genes in the same cell resulted in gRNA depletion. Further, we found that the level of depletion from the screen for all titration pairs correlates with the predicted activity for partial knockdown gRNAs (**Fig. 2c, Supplementary Fig. 4a-d**).

To validate that the lack of collateral activity in this screen is a result of low copy number of *Rfx*Cas13d, we performed a competition assay and targeted *HDAC1* and/or *HDAC2* using either transfected or transduced *Rfx*Cas13d (**Fig. 2d, e**). As before, we observed depletion when both *HDAC1* and *HDAC2* were targeted, regardless of how *Rfx*Cas13d was delivered. However, when targeting *HDAC1* alone, we only observed depletion of cells when *Rfx*Cas13d was delivered via transfection. Notably, targeting *HDAC1* using perfect match gRNAs resulted in a significant decrease in GFP in the transfection condition. Even with imperfect match gRNAs (predicted knockdown of 55%), we observed a slight decrease in GFP signal in the transfection condition. This confirms that modulating the rate of target recognition with designed mismatches in the gRNA not only alters *cis* RNA cleavage but also *trans* collateral cleavage.

Due to increasing interest in high-fidelity variants of *Rfx*Cas13d for transcriptome engineering^16,17^, we wanted to evaluate the activity of a recently described high-fidelity variant (hfCas13d)^8^ using low-copy number lentiviral delivery. We generated a doxycycline-inducible hfCas13d HAP1 cell line via lentiviral transduction and, using it, targeted 4,890 genes with eight gRNAs each to compare the nuclease activity of hfCas13d to *Rfx*Cas13d^13^ (**Fig. 3a, Supplementary Table 4**). We first validated the cell line using gRNAs targeting the surface markers CD46 and CD55 and found that at low copy number, hfCas13d has a ∼4-fold weaker knockdown efficiency even though nuclease expression is similar between the two cell lines (**Supplementary Fig. 5a**). For the top 5 most depleted genes from the hfCas13d or *Rfx*Cas13d pooled screens, we see weaker gRNA depletion with hfCas13d and greater variation between gRNAs targeting the same gene (**Fig. 3b**).

Notably, while some gRNAs did deplete from the hfCas13d screen, the vast majority of genes, even those that are highly essential, did not — a stark contrast to *Rfx*Cas13d (**Fig. 3c, Supplementary Fig. 5b, c**). When comparing *Rfx*Cas13d and hfCas13d screens on a gene level, we found that 1,235 genes have a 2-fold or greater depletion in the *Rfx*Cas13d screen, whereas only 2 of the 4,890 genes (*SNRPD3* and *SMU1*) have a 2-fold or greater depletion in the hfCas13d screen (**Fig. 3d, Supplementary Table 5**). For high-confidence essential genes, we identified 94% of these genes depleted in the *Rfx*Cas13d screen but only 3% depleted in the hfCas13d screen (DepMap Chronos essentiality score ≤ -1, *n* = 785 genes) (**Fig. 3e**). Importantly, we successfully discriminated essential genes from nonessential genes with *Rfx*Cas13d (AUROC: 0.96), but not with hfCas13d (AUROC: 0.62), due to a lack of essential gene depletion (**Fig. 3f**).

These results indicate that the regulation of *Rfx*Cas13d expression is important to mitigate risks of collateral RNA degradation. When using lentiviral transduction at low copy, we find that *Rfx*Cas13d is capable of high on-target transcript knockdown with no indication of collateral RNA degradation. While high-fidelity Cas13 variants can reduce collateral activity risks, it comes at the expense of lower on-target knockdown. Given the rapid advancement of transcriptome engineering and forward transcriptomic screens using Cas13, careful consideration of Cas13 expression and delivery methods is necessary to ensure high on-target activity and minimal collateral RNA degradation.

## Acknowledgements

We thank the entire Sanjana laboratory for support and advice. We thank the New York University Biology Genomics Core for sequencing resources. N.E.S. is supported by New York University and New York Genome Center funds, National Institutes of Health (NIH)/National Human Genome Research Institute (DP2HG010099, R01HG012790), NIH/National Cancer Institute (R01CA218668, R01CA279135, R21CA272345), NIH/National Institute for Allergy and Infectious Diseases (R01AI176601), the MacMillan Center for the Study of the Noncoding Cancer Genome at the New York Genome Center, and the Simons Foundation for Autism Research (Genomics of ASD 896724).

## Author contributions

S.K.H. and N.E.S conceived the project. H-H.W. and N.E.S. designed the Cas13 libraries. S.K.H., H-H. W., and A.M-M. performed pooled Cas13 screens. S.K.H., S.M., G.D., and O.C. conducted validation assays. S.K.H., H-H.W. and S.M. analyzed pooled screens. N.E.S. supervised the work. S.K.H. and N.E.S. wrote the manuscript with input from all authors.

## Additional information

Correspondence and requests for materials should be addressed to N.E.S.

## Methods

### Cloning of RfxCas13d-eGFP and high-fidelity Cas13d (hfCas13d) vectors

We modified pLentiRNACRISPR_007 (Addgene 138149, TetO-NLS-*Rfx*Cas13d-NLS-WPRE-EFS-rtTA3-2A-Blast) by replacing the blasticidin resistance gene with GFP via Gibson cloning. We termed this vector pLentiRNACRISPR_009. We also modified the pLentiRNACRISPR_007 and replaced *Rfx*Cas13d with hfCas13d N2V8^8^ using Gibson cloning. hfCas13 is a modified form of *Rfx*Cas13d with the following point mutations: A134V, A140V, A141V, A143V. We termed this vector pLentiRNACRISPR_008. All constructs were confirmed by Sanger and nanopore (full-plasmid) sequencing (Azenta and plasmidsaurus). All primers used for molecular cloning and guide sequences are shown in **Supplementary Table 6**.

### Monoclonal Cas13 cell line generation and cell culture

HEK293FT and A375 cells were cultured in D10 medium, which is Dulbecco’s Modified Eagle Medium (DMEM) with high glucose and stabilized L-glutamine (Caisson DML23) supplemented with 10% fetal bovine serum (Sigma 14009C). HAP1 cells were cultured in I10 medium: Iscove’s Modified Dulbecco’s Medium (IMDM) with L-glutamine (Caisson IML02) supplemented with 10% fetal bovine serum. All cells were incubated at 37 °C with 5% carbon dioxide.

Monoclonal doxycycline-inducible hfCas13d-NLS HAP1 cells were generated by transducing cells with an hfCas13d-expressing lentivirus at a low multiplicity of infection (< 0.1) and selected with 5 μg/ml of blasticidin S (AG Scientific B-1247). Single-cell colonies were isolated by low-density plating and then expression of HA-tagged hfCas13 was confirmed by immunoblot using an anti-HA peptide antibody (Cell Signaling Technology 2367S). hfCas13 knockdown activity was confirmed by flow cytometry using APC anti-CD46 and FITC anti-CD55 (BioLegend 352405 and 311306, respectively). Monoclonal doxycycline-inducible *Rfx*Cas13d-NLS HEK293FT, A375 and HAP1 cells were previously validated by Wessels and Méndez-Mancilla et al^4^ and Guo et al^11^. All monoclonal cell lines with Cas13 (or variants) were cultured with 5 μg/ml of blasticidin S.

### Plasmid transfection of RfxCas13d

Plasmids were transfected using Lipofectamine 3000 (Invitrogen L3000008). HEK293FT cells were seeded at a density of 2 × 10^5^ cells per well in a 12-well plate and transfected the next day. For each well, 3 uL of Lipofectamine 3000 reagent was added to 50uL of prewarmed Opti-MEM I Reduced-Serum Medium (Gibco 31985062) and subsequently mixed with 50uL Opti-MEM containing 2uL P3000 reagent, 250 ng of *Rfx*Cas13d or pUC19 vector, and 50 ng of pmaxGFP (Lonza). Following a 10-minute incubation at room temperature, the mixture was added dropwise to cells. At 48 hours post-transfection, we performed quantitative imaging or harvested cells for flow cytometry or reverse transcription followed by quantitative PCR. Of relevance to the transfection experiments, the Cas13 (lentiviral) plasmids contain a CMV promoter before the 5’ LTR.

### Lentiviral production and transduction of guide RNA vectors

Lentivirus was produced via transfection of plasmids expressing *Rfx*Cas13d guide RNAs under the U6 promoter with lentiviral packaging plasmids using linear polyethylenimine (PEI) MW25000 (Polysciences 23966) in HEK293FT. Cells were seeded at one million per well in a 6-well plate and transfected with 7.5 μl PEI, 1.2 μg gRNA plasmid, 0.825 μg psPAX2 (Addgene 12260) and 0.55 μg pMD2.G (Addgene 12259). Two days post-transfection, the viral supernatant was collected and filtered through a 0.45-μm filter and then stored at −80 °C until further use. For transductions of single or dual guide RNAs, 200uL of lentivirus was added to 250,000 cells of the appropriate cell line along with 8ug/mL polybrene (Santa Cruz Biotechnology, sc-134220). The media was changed 24 hours after transduction with 1 μg/ml puromycin (Thermo A1113803) added. Puromycin selection was completed within 48 hours for all cell lines, as determined by complete killing of untransduced control cells in puromycin media.

### Western blot

Transduced or transfected HEK293FT cells were collected 48h after doxycycline-induction (1 μg/ml), washed with 1x PBS and lysed with total lysis buffer (50 mM Tris (pH 7.4), 50 mM NaCl, 2 mM MgCl2 and 1% SDS) supplemented with Benzonase (MilliporeSigma 70746-4, 0.5 μl per 100 μl lysis buffer) for 15 minutes at room temperature. The protein concentration was determined using the BCA protein assay (Thermo 23227). Equal amounts of cell lysates (20 μg) were denatured in NuPAGE LDS Sample Buffer (Thermo P0007) supplemented with 100 mM DTT for 10 minutes at 65°C. Denatured samples and PageRuler pre-stained protein ladder (Thermo 26616) were separated in Novex 4-12% Tris-Glycine mini gels (Thermo XP04125) in 1x Tris-Glycine-SDS buffer (IBI Scientific IBI01160) for 1 hour at 180 V. Proteins were transferred on a nitrocellulose membrane (BioRad 1620112) in 1x Tris-Glycine transfer buffer (Thermo LC3675) supplemented with 10% methanol for 1 hour at 30 V. Immunoblots were blocked with 5% BSA (VWR AAJ65097) dissolved in in 1x TBS with 1% Tween-20 (TBS-T) for 60 min and separately incubated with primary antibodies: anti-HA (Cell Signaling Technology 2367S, 1:2000 dilution in 5% BSA/TBS-T) and anti-beta-tubulin (Invitrogen 32-2600, 1:2000 dilution in 5% BSA/TBS-T) overnight at 4°C. Following the primary antibody, immunoblots were incubated with IRDye 800CW donkey anti-mouse (LI-COR 925-32212, 1:10,000 dilution in 5% BSA/TBS-T). The blots were imaged using an Odyssey CLx (LI-COR).

### Flow cytometry

For flow cytometry analysis, 2 × 10^5^ cells per condition were stained with CD46 antibody (BioLegend 352405: anti-CD46 clone TRA-2-10, 0.7 μl per 2 × 10^5^ cells). Cells were gated by forward and side scatter and signal intensity to remove potential multiplets and additionally gated for living cells using DAPI exclusion (ThermoFisher L34963). For each sample we analyzed at least 5,000 cells. If cell numbers varied, we downsampled all conditions to the same number of cells before calculating the mean fluorescence intensity.

### Quantitative microscopy

GFP was imaged using a BZ-X Filter FITC (Keyence OP-87764) on a BZ-X810 microscope (Keyence) or with a Incucyte S3 (Sartorius). Images for measurement were taken at 10× magnification. For quantification using the Keyence microscope, 12 representative images were taken in each well and the mean fluorescence intensity (per image) was averaged across all images to generate a mean fluorescence intensity per well.

For live cell imaging of HEK293FT cells transfected with differing amounts of pLentiRNACRISPR_009, cells were first transduced with lentivirus containing *MIF* or non-targeting gRNAs and selected with 1 μg/ml puromycin for 48 hours. They were then transfected with differing amounts of pLentiRNACRISPR_009 and 9 representative images were taken from each well using a Incucyte S3. The GFP mean fluorescence intensity was averaged across all images to generate the mean fluorescence intensity per well over the course of 2 days following transfection. Each well was first normalized by the intensity of the initial time point post-Cas13d transfection. Then, collateral loss of GFP was calculated by dividing the mean fluorescence intensity per well for cells transduced with *MIF* gRNAs by those transduced with non-targeting gRNAs over the time course.

### Competition assays for cell growth and live imaging

For the competitive cell growth assays with GFP readout, we exchanged the PuroR cassette in pLentiRNAGuide_001 (Addgene 138150) with a GFP-P2A-Puro cassette from pLentiRNAGuide_003 (Addgene 192505) using *NheI* and *ApaI* restriction sites. We termed this new vector pLentiRNAGuide_004. We transduced A375-*Rfx*Cas13d or A375 wild-type cells with lentivirus containing dual array gRNAs in pLentiRNAGuide_004 vector. We performed independent transductions for each gRNA array in triplicate and selected cells with 1 ug/mL puromycin for 3 days. The GFP-positive cells were co-cultured with parental cells in equal ratios for 24 hours and the ratio of GFP-positive to total cells was determined using a Incucyte S3 (Sartorius). Cas13 expression was then induced with 1ug/mL doxycycline and the GFP-positive/GFP-negative cell ratio was observed over the course of 5-7 days with 9 images per well being taken at 8-hour intervals. Survival of perturbed cells was calculated by normalizing ratios to the initial time point pre-Cas13 induction per well and comparing to the median of cell mixtures containing cells that were transduced with non-targeting gRNA arrays. Representative images show confluence masks of GFP-positive and GFP-negative cells. All gRNA sequences for the arrayed validation are given in **Supplementary Table 6**.

### Quantitative reverse transcription quantitative PCR (RT-qPCR)

RNA was isolated using the Direct-zol RNA Purification Kit (Zymo R2062). Synthesis of cDNA was performed using RevertAid Reverse Transcriptase (Thermo EP0442). Quantitative RT-qPCR was performed using a QuantStudio 5 instrument (Applied Biosystems) with Luna Universal qPCR Master Mix (NEB M3003E). Relative transcript abundance was normalized to *GAPDH* and non-targeting control gRNAs (ΔΔCt method). Primer sequences can be found in **Supplementary Table 6**.

### Pooled Cas13 libraries: Design and cloning

In total, we performed 3 pooled Cas13 screens (Screen 1: nonessential and essential genes; Screen 2: HDAC1 and HDAC2 paired titration; Screen 3: high-fidelity Cas13 screen) and the design and cloning of each library is described below.

For Screen 1, we designed a *Rfx*Cas13d gRNA library targeting 4,177 human protein-coding gene transcripts. The genes were categorized into 3 different groups of essentiality based on all CRISPR-Cas9 screens from Hart et al.^14^ with 3,448 genes being nonessential (DepMap release 05-2023; -0.25 ≤ DepMap score ≤ 0.5), 488 selectively essential (−1 ≤ DepMap score <-0.25), and 241 essential (DepMap score < - 1). To rationally identify a relationship between target gene expression and collateral RNA degradation, we chose genes over a large range of gene expression values (spanning <1 to >1000 transcripts per million reads).

For this pooled screen, we designed optimized gRNAs using a model for Cas13 gRNA design trained on thousands of gRNAs (http://cas13design.nygenome.org)^11^. For each transcript, we selected 5 gRNAs from the highest (or second-highest as needed) efficacy quartile (as given by cas13design). We also embedded 1,000 non-targeting gRNAs as negative controls, which we ensured had 3 or more mismatches to any other transcripts (hg19). In total, the library included 17,708 gRNAs. Each gRNA was flanked with constant regions (for PCR amplification and Gibson cloning) and synthesized as 106mer single-stranded oligonucleotides (Twist Biosciences). A full list of gRNA sequences for the Screen 1 library can be found in **Supplementary Table 1**.

For Screen 2 (*HDAC1* and *HDAC2* synthetic lethality titration screen), we first selected 20 pairs of target sites for each gene. For each target site (in *HDAC1* or *HDAC2*), we designed 1 perfect-match gRNA (i.e. the target site itself) and 10 additional single-mismatch variant gRNAs with titrated transcript knockdown activities ranging from 5-95% (incrementing at every decile) that were designed using the TIGER model library (10). Each target site in *HDAC1* was paired with each target site in *HDAC2* in a dual gRNA array. For each target site pair, we used 11 *HDAC1* titration gRNAs × 11 *HDAC2* titration gRNAs × 2 positions (*HDAC1*-*HDAC2* gRNA order or *HDAC2*-*HDAC1* gRNA order), which yielded 242 combinations for each target site pair. Given the 20 pairs of target sites, we generated a total of 4,839 *HDAC1*/*HDAC2* arrays. We also paired each gRNA targeting *HDAC1* or *HDAC2* with a non-targeting (NT) gRNA: This yielded an additional 880 arrays = 11 gRNAs per target site x 40 total target sites (20 in *HDAC1*, 20 in *HDAC2*) × 2 positions (*HDAC1*/*2-*NT gRNA order or NT*-HDAC1/2* gRNA order). We also included 492 arrays with NT gRNAs in both positions. In total, the library for Screen 2 consisted of 6,211 arrays. Each gRNA was flanked with constant regions (for PCR amplification and Gibson cloning) and synthesized as 148mer single-stranded oligonucleotides (Twist Biosciences). A full list of gRNA sequences for the Screen 2 library can be found in **Supplementary Table 3**.

For Screen 3 (hfCas13 screen), we utilized one of the libraries from prior work used to train the TIGER deep learning model^13^. Briefly, the library contained gRNAs targeting 4,890 genes of varying degrees of essentiality as follows: 785 highly essential genes (DepMap release 05-2023; DepMap score < −1 in ≥1000 cell lines), 722 genes with medium essentiality (−1 ≤ DepMap score <-0.5), 1,398 genes with low essentiality (−0.5 ≤ DepMap score <-.5), and 1,985 nonessential genes (DepMap score ≥ -0.1), all spanning a wide range of gene expression values from 1 to >1,000 TPM. For all 4,890 target genes, we designed eight gRNAs targeting the genes protein-coding region. We added 998 non-targeting control gRNAs with more than three mismatches to the hg19 transcriptome. In total, the library consisted of 40,118 gRNAs. Each gRNA was flanked with constant regions (for PCR amplification and Gibson cloning) and synthesized as 106mer single-stranded oligonucleotides (Twist Biosciences). A full list of gRNA sequences for the Screen 3 library can be found in **Supplementary Table 4**.

For library cloning, crRNA libraries were amplified using 8 replicate 50-ul PCR reactions with 8 amplification cycles. Following gel-purification, the resulting amplicon was Gibson-cloned into BsmBI-digested pLenti*Rfx*Guide-Puro (Addgene 138151). We verified successful cloning via Illumina sequencing to verify high gRNA recovery (>99%) and minimal bias (90:10 ratio < 5).

### Pooled lentiviral production

Lentivirus was produced via transfection of pooled library plasmids with appropriate packaging plasmids (psPAX2-Addgene 12260; pMD2.G-Addgene 12259) using linear polyethylenimine (PEI) MW25000 (Polysciences 23966) in HEK293FT. Cells were seeded at ten million per 10 cm dish and transfected with 60 μl PEI, 9.2 μg transfer plasmid pool, 6.4 μg psPAX2 and 4.4 μg pMD2.G. Three days post-transfection, the viral supernatant was collected and filtered through a 0.45-μm filter and then stored at −80 °C until further use. To minimize the introduction of multiple gRNAs per cell, the lentivirus volume used for transduction was titrated to result in 30 - 40% survival after puromycin selection.

### Pooled Cas13 library CRISPR screens

Cas13d-expressing cells were transduced with the pooled library lentivirus, ensuring at least a 1000x guide representation in the pool of cells per separate infection replicate. They were transduced via spinfection at 1000 rpm for 1 hour at 30 °C (Beckman Coulter, Allegra X-14R), followed by overnight incubation. After 24 hours, media was changed and supplemented with 1 μg/ml puromycin (Thermo A1113803) added. Puromycin selection was completed within 48 hours for all cell lines.

Following puromycin selection, Cas13d expression was induced by replenishing with growth medium containing 1 μg/ml puromycin, 5 μg/ml blasticidin and 1 μg/ml doxycycline. Cells were passaged every 2 to 4 days and split as needed. Samples were harvested at 0, 7, and 14 days post-Cas13d induction. Genomic DNA was isolated from cell pellets with a representation of at least 1,000 cells per construct using the following protocol^18^: for 100 million cells, 12 ml of NK lysis buffer (50 mM Tris, 50 mM EDTA, 1% SDS and pH 8) was used to lyse cell pellets. 60 μl of 20 mg/ml Proteinase K (Qiagen) was added to resuspended cells and incubated at 55 °C overnight. The next day, 60 μl of 20 mg/ml RNase A (Qiagen) was added to cells and incubated at 37 °C for 30 min. After, 4 ml of prechilled 7.5 M ammonium acetate was added and samples were vortexed and spun at 4,000g for 10 min. The supernatant was moved to a new tube and mixed well with 12 ml isopropanol before being spun at 4,000g for 10 min. DNA pellets were washed with 12 ml of 70% ethanol and then spun down. After ethanol removal, pellets were dried and then resuspended with 0.2× TE buffer (Sigma-Aldrich).

We amplified gRNA cassettes from the genomic DNA using a two-step PCR protocol (PCR1 and PCR2). PCR1 was performed to amplify a region containing the crRNA cassette in the lentiviral genomic integrant using TaqB polymerase (Enzymatics P7250L). We performed an appropriate amount of PCR1 reactions for each gDNA sample based on library size (Screen 1: 40 reactions; Screen 2: 20 reactions; Screen 3: 48 reactions) using 10 μg gDNA per 100 μl PCR1 reaction (10 cycles). We then combined PCR1 products for the same sample together before PCR2, which was done to incorporate Illumina adaptors using Q5 polymerase (NEB M0491). We performed an appropriate number of PCR2 reactions for each sample (Screen 1: 16 reactions; Screen 2: 8 reactions; Screen 3: 24 reactions) using 5 μl unpurified PCR1 product per 50 μl reaction (18 cycles). The resulting amplicons from PCR2 (∼270 bp) were pooled and then purified using SPRI beads (Beckman B23317) or gel extracted using a QiaQuick Gel Extraction kit (Qiagen 28704). The concentration of the purified PCR product was quantified using Qubit dsDNA HS Assay Kit (Thermo Q32851) and sequenced on an Illumina NextSeq 500 using a single ended 150 cycle read.

### Pooled screen analysis

We processed reads from pooled Cas13d screens following established pipelines^4^: First, reads were de-multiplexed based on Illumina i7 barcodes present in PCR2 reverse primers and custom in-read i5 barcodes, allowing for one mismatch. Reads were trimmed to the expected gRNA length by identifying known anchor sequences relative to the guide sequence. Reads were trimmed using Cutadapt (v.1.13)^19^ with the following parameters: `-g CTGGTCGGGGTTTGAAAC -e 0.2 -O 5 --discard-untrimmed` and -`a TTTTTGAATTCGCTAGCT -e 0.1 -O 5 --minimum-length 15 --discard-untrimmed`.

We aligned processed reads to the designed crRNA reference using bowtie (v.1.1.2)^20^ and allowed for up to three mismatches (parameters: `-v 1 -m 3 –best -q`) for Screen 1 and Screen 3. For Screen 2 (*HDAC1* and *HDAC2* titration), processed reads were collapsed (*FASTX-Toolkit*) to count perfect duplicates followed by exact string-match intersection with the reference such that only perfectly matching and unique alignments were kept. The raw gRNA counts were normalized using a median of ratio method^21^ and batch corrected for biological replicates using combat from the SVA R package (v.3.34.0)^22^. Nonreproducible technical outliers were removed by pair-wise linear regression for each sample, collecting residuals, and taking the median value for individual gRNAs across each time point.

To calculate gRNA depletion, count ratios between the desired time point (Day 7 or Day 14) and the early timepoint (Day 0) sample for each replicate were computed followed by log2 transformations. Consistency across replicates was assessed using Pearson correlations and robust rank aggregation^23^. For Screen 2 (HDAC1/2 titration), which contains gRNAs with mismatches, we computed the ratio of the fold-change of the permuted gRNA to the fold-change of the perfect match reference gRNA^13^; these computations were performed in log space.

