## Supplementary Figures for "Low copy CRISPR-Cas13d mitigates collateral RNA cleavage"

**Supplementary Tables**

**Supplementary Table 1.** Collateral activity screen target gRNA depletion.

**Supplementary Table 2.** Collateral activity screen gene depletion.

**Supplementary Table 3.** HDAC1/HDAC2 titration screen gRNA annotation and depletion.

**Supplementary Table 4.** High-fidelity Cas13 screen gRNA depletion.

**Supplementary Table 5.** High-fidelity and wild-type Cas13 screen gene depletion.

**Supplementary Table 6.** Oligonucleotides used in this study.

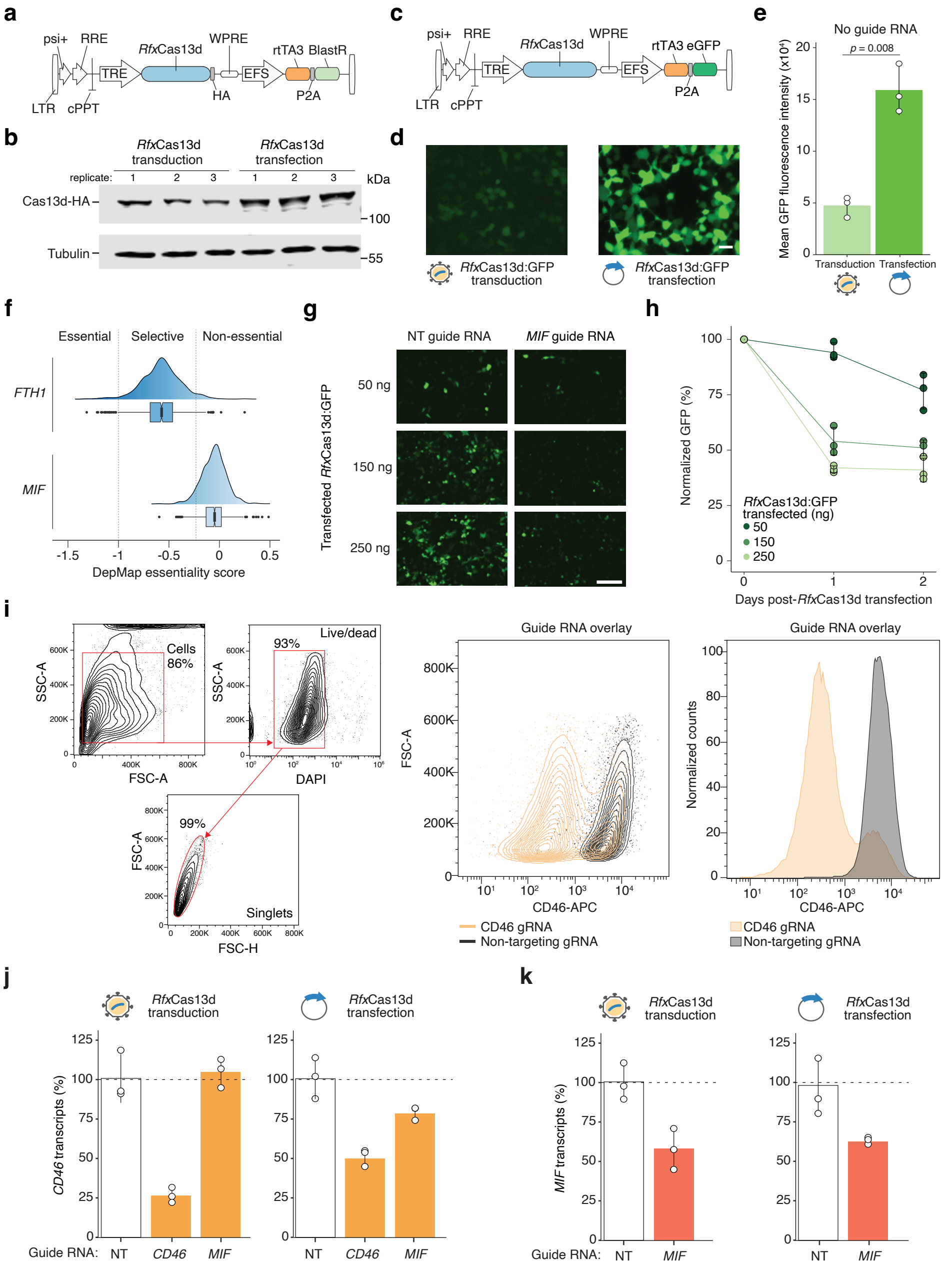


**Supplementary Figure 1. *Rfx*Cas13 expression and individual guide RNA flow gating and on-target knockdown.**

**a,** Doxycycline-inducible *Rfx*Cas13d that co-expresses blasticidin resistance (pLentiRNACRISPR_007).

**b,** Western blots of HA-tagged *Rfx*Cas13d and beta-tubulin expression where *Rfx*Cas13d was either transduced or transfected in HEK293FT cells ($n$ = 3 biological replicates).

**c,** Doxycycline-inducible *Rfx*Cas13d that co-expresses EGFP (pLentiRNACRISPR_009).

**d, e,** GFP imaging (*d*) and quantification (*e*) of HEK293FT cells 48 hours after being either transfected or transduced with pLentiRNACRISPR_009 without any guide RNA present. Scale bar: 50 µm. Significance was determined using a two-sided *t*-test ($n = 3$ biological replicates).

**f,** DepMap essentiality score density plot for *FTH1* and *MIF* across 1,095 cell lines. Dashed lines indicate essentiality group (Essential: mean DepMap Chronos score [DMCS] $< -1$, Selective: $-1 \leq$ DMCS $<-0.25$ , and nonessential: DMCS $> -0.25$).

**g, h,** GFP imaging (*g*) and quantification (*h*) of HEK293FT cells previously transduced with *MIF*-targeting or non-targeting (NT) gRNAs 48 hours after being transfected with 50, 150, or 250 nanograms of pLentiRNACRISPR_009. GFP expression was normalized to non-targeting control gRNAs ($n = 3$ biological replicates). Scale bar: 200 µm.

**i,** Representative flow cytometry gating for the readout of CD46 expression (collateral degradation from *Fig. 1d, e*).

**j, k,** *CD46* (*j*) and *MIF* (*k*) RNA knockdown using gRNAs from *Fig. 1c, e* where Cas13d was either transduced or transfected. RNA knockdown is measured by RT-qPCR relative to cells transduced with a non-targeting gRNA as a control and was normalized to *GAPDH*.

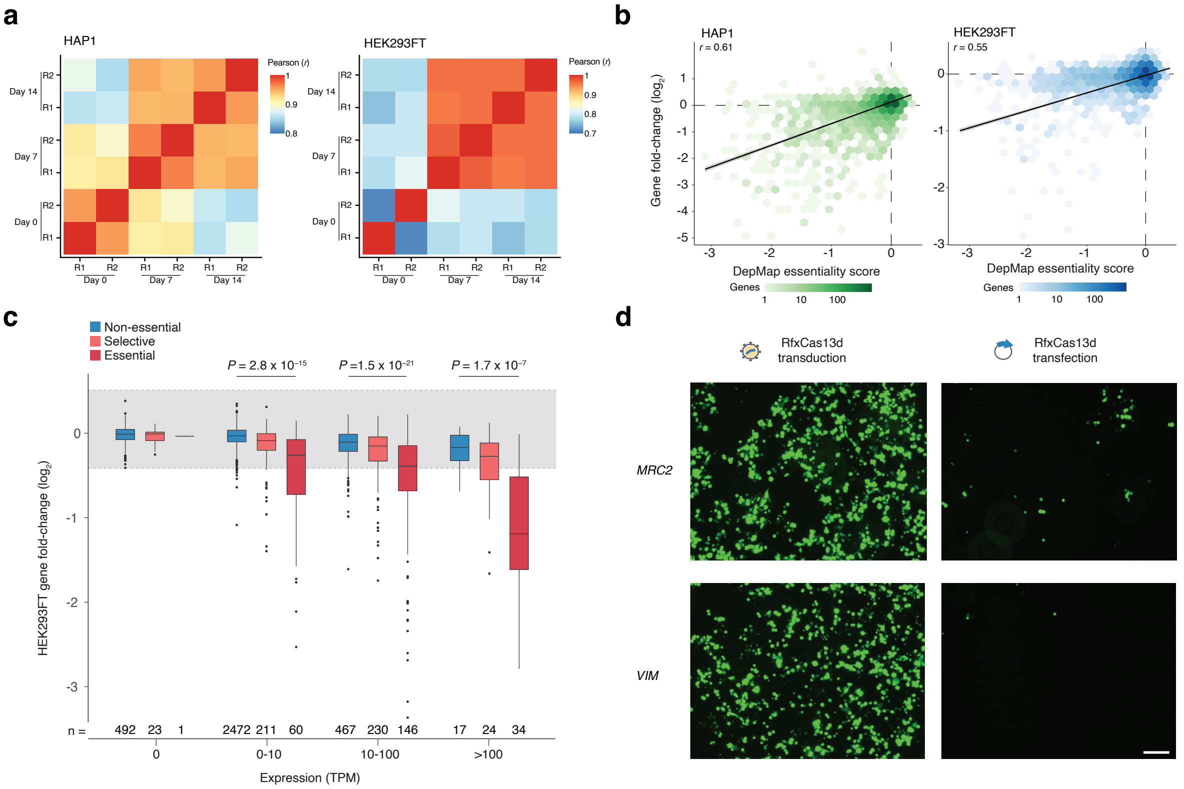


**Supplementary Figure 2. *Rfx*Cas13d pooled screen replicate correlations and comparisons to DNA-targeting (Cas9) pooled screens.**

**a,** Pearson (*r*) correlation coefficients of gRNA counts across both replicates for the screen performed in *Fig. 1f* in both HAP1 and HEK293FT cell lines.

**b,** Correlation between fold change of the 4,177 genes from the Cas13d screens in HAP1 and HEK293FT and DepMap scores from Cas9 perturbation studies[14]. Bin colors represent gene counts.

**c,** Fold-change of genes compared to their expression from the HEK293FT Cas13d screen, grouped by their essentiality status defined by their mean DepMap Chronos score (DMCS): essential: DMCS < -1, selective: -1 ≤ DMCS < -0.25, non-essential: DMCS ≥ -0.25, *n* = 1,095 cell lines). Statistical significance between non-essential and essential gene categories was determined by a two-sided Mann-Whitney *U* test.

**d,** Representative images of HEK293FT cells transduced or transfected with Cas13d with gRNAs targeting the nonessential genes *MRC2* and *VIM*. Scale bar: 100 µm.

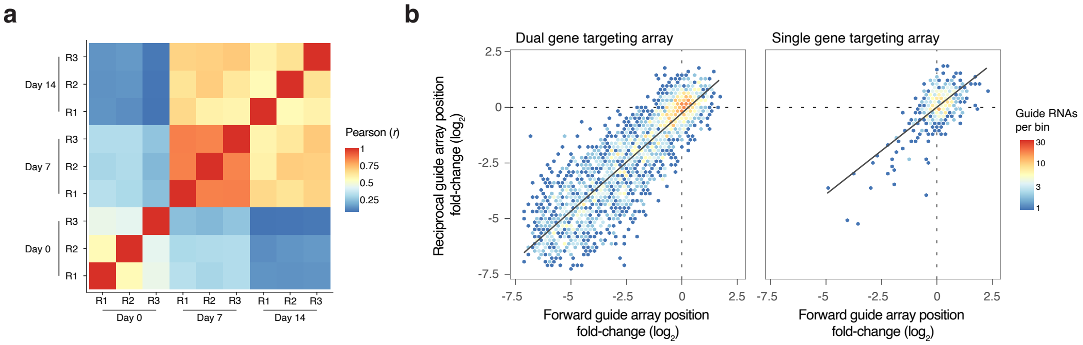


**Supplementary Figure 3. HDAC1/HDAC2 synthetic lethality titration pooled screen replicate correlations and quality control of dual guide RNA arrays.**

**a,** Pearson (*r*) correlation coefficients of gRNA counts across 3 replicates for the screen performed in *Fig. 2a* in A375 cells.

**b,** Fold change of all gRNA arrays in the *HDAC1/HDAC2* synthetic lethality titration screen comparing the forward guide array position (gRNA-1,gRNA-2) versus its reciprocal position (gRNA-2,gRNA-1) when both *HDAC1* and *HDAC2* are targeted (dual gene targeting array, *left*; Pearson *r* =0.88) or when either *HDAC1* or *HDAC2* are targeted along with a non-targeting gRNA (single gene targeting array, *right*; Pearson *r* =-0.09).

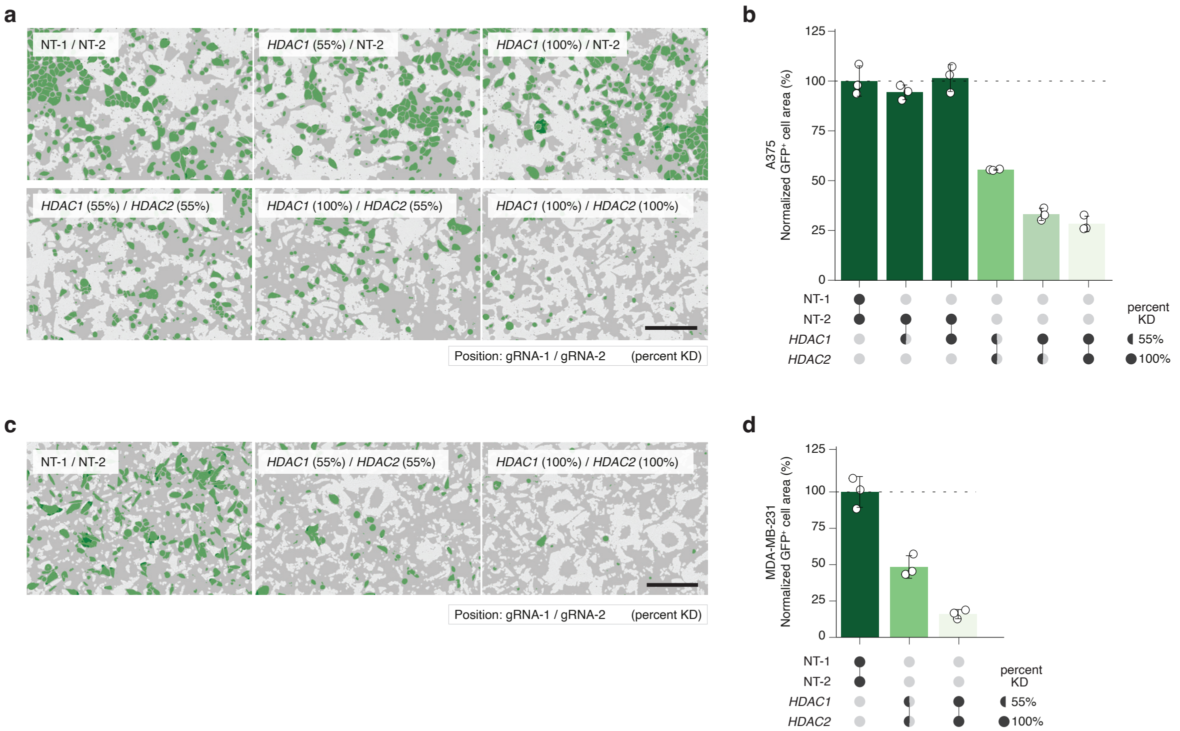


**Supplementary Figure 4. Arrayed validation of HDAC1/HDAC2 synthetic lethality titration in human A375 and MDA-MB-231 cells.**

**a,** Representative images of A375 cells transduced with Cas13d with dual gRNA arrays targeting *HDAC1* and *HDAC2* at 55% or 100% predicted gRNA activity or non-targeting gRNAs. Scale bar: 200 µm.

**b,** Percent of GFP-positive A375 cells transduced with Cas13d for each gRNA array normalized to non-targeting gRNA array control ($n$ = 27 images from 3 independent transductions).

**c,** Representative images of MDA-MB-231 cells transduced with Cas13d with dual gRNA arrays targeting *HDAC1* and *HDAC2* at 55% or 100% predicted gRNA activity or non-targeting gRNAs. Scale bar: 200 µm.

**d,** Percent of GFP-positive MDA-MB-231 cells transduced with Cas13d for each gRNA array normalized to non-targeting gRNA array control ($n$ = 27 images from 3 independent transductions).

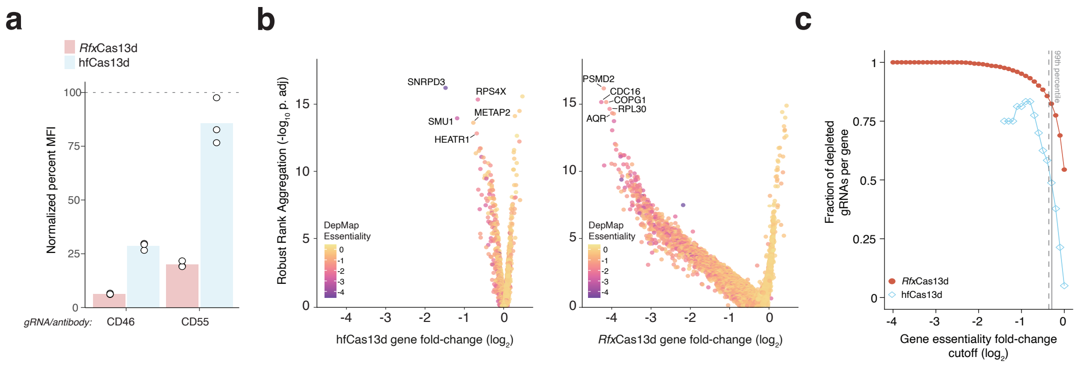


**Supplementary Figure 5. *Rfx*Cas13d and hfCas13d on-target knockdown and pooled screen essential gene depletion.**

**a,** On-target knockdown of CD46 and CD55 by flow cytometry in the *Rfx*Cas13d and hfCas13d HAP1 cell lines. Cells were transduced with gRNAs targeting *CD46* or *CD55* and normalized to cells that were transduced with a non-targeting gRNA.

**b,** Depletion of essential and nonessential genes in hfCas13d and *Rfx*Cas13d HAP1 cell lines by collapsing individual gRNAs into corresponding genes using Robust Rank Aggregation (RRA). Gene essentiality is defined by its DepMap Chronos score.

**c,** Fraction of active gRNAs (out of 8 gRNAs per gene) as a function of gene depletion. Gray lines indicate the 99$^{th}$ percentile of NT gRNA distribution for *Rfx*Cas13d (*solid*) and hfCas13d (*dashed*) screens.
